# Shift in vacuolar to cytosolic regime of infecting *Salmonella* from a dual proteome perspective

**DOI:** 10.1101/2023.02.07.527450

**Authors:** Ursula Fels, Patrick Willems, Margaux De Meyer, Kris Gevaert, Petra Van Damme

## Abstract

By applying dual proteome profiling to *Salmonella enterica* serovar Typhimurium (*S*. Typhimurium) encounters with its epithelial host (here, *S*. Typhimurium infected human HeLa cells), a detailed interdependent and holistic proteomic perspective on host-pathogen interactions over a time course of infection was obtained. Data-independent acquisition (DIA)-based proteomics was found to outperform data-dependent acquisition (DDA) workflows, especially in identifying the downregulated bacterial proteome response during infection progression infection by permitting quantification of low abundant bacterial proteins at early times of infection at low bacterial infection load. *S*. Typhimurium invasion and replication specific proteomic signatures in epithelial cells revealed interdependent host/pathogen specific responses besides pointing to putative novel infection markers and signalling responses.

## Introduction

*Salmonella enterica* serovar Typhimurium (*S*. Typhimurium) is an enteric facultative intracellular pathogen with wide-ranging disease outcomes grading from self-limiting gastroenteritis to systemic infection. As a foodborne pathogen, *S*. Typhimurium invades the intestinal tissue via different routes through infection of a diverse range of host cells, including epithelial cells (e.g., M cells and absorptive epithelial cells) and immune cells (e.g., dendritic cells and macrophages), in which they proliferate (1, 2). Infection by *Salmonella* is mediated by its two type III secretion systems (T3SSs); T3SS-1 and −2, encoded within *Salmonella* pathogenicity island 1 (SPI-1) and −2 (SPI-2), respectively (3). Along with the T3SS-1 components, the 40-kb SPI-1 gene cluster encodes chaperones, type III effector proteins (T3Es) and transcriptional regulators that modulate expression of many virulence genes within, but also outside SPI-1 (4). Whereas T3SS-1 and its effectors are mainly associated with the invasion of non-phagocytic epithelial cells (5), T3SS-2 delivers effectors that promote *Salmonella* replication inside phagocytic as well as non-phagocytic cells (6–8). To invade epithelial cells, *Salmonella* delivers through its T3SS-1 a first batch of effectors into the host cytoplasm to provoke actin cytoskeleton rearrangements that allow for bacterial uptake by macropinocytosis. More specifically, lamellipodia and filopodia-containing protrusions, called membrane ruffles, emerge and enclose the pathogen (9, 10). This active invasion of epithelial cells mediated by T3SS-1 and its cognate effectors is thought to play a key role in gut inflammation during salmonellosis (1, 5). A first set of pro-inflammatory responses (e.g., NF-κB signalling) are detected upon entry of the pathogen owed to both manipulation of cell death pathways and/or tight junctions via effectors, and by recognition of peptidoglycan, flagellin or lipopolysaccharide by host pattern recognition receptors (PRRs) (11). Following internalization, *Salmonella* resides inside a phagosome-like compartment, dubbed the *Salmonella*-containing vacuole (SCV), where it further rewires host cell processes to create a replicative niche mainly through the action of T3SS-2 effectors (12). In essence, T3SS-2 effectors maintain the SCV membrane, navigate juxtanuclear SCV positioning and elicit filamentous membrane extensions from the SCV along microtubules, termed *Salmonella*-induced filaments (SIFs) (13). SIFs have been observed in various cell types and exhibit endosomal markers like LAMP1, Rab7 and vacuolar ATPase (13). Interestingly, the onset of SIF biogenesis at 4-8 hours post-infection coincides with intercellular bacterial replication and has therefore been postulated to facilitate nutrient acquisition from the host, amongst others (14–17). Remarkably, in about 20% of infected epithelial cells, bacterial replication also occurs outside the SCV in the cytosol where flagellated bacteria hyper-replicate and account for the majority of net replication (18). Moreover, some SCVs are displaced from their juxtanuclear positioning in the host cell towards the cell periphery (19). Both phenomena, i.e., vacuolar escape and centrifugal SCV movement, have been linked to dissemination of the pathogen to other cells to repeat the infectious cycle (19). Moreover, the various (subcellular) environments encountered by these bacterial subpopulations during infection further lie at the basis of the observed heterogeneity of bacterial subpopulations *in vitro* (20) and *in vivo* (21). Specific bacterial subpopulations were for example found to reprogram infected hosts to promote long-term bacterial survival (22).

As such, it is clear that over the course of an infection, an intricate and complex interplay between *Salmonella* and its host occurs that ultimately defines the infection outcome. As diverse successive environmental cues shape both the host and bacterial proteome, this highlights the importance of studying both proteomes in an integrative manner to unwire this multifaceted interaction and to obtain holistic insights. The characterization of molecular mechanisms underlying interactions between pathogen and host has extensively advanced over the last two decades, mainly through omics-informed infection studies in cellular as well as animal infection models.

Staples *et al*. (22) characterized non-growing, metabolically active subsets of *Salmonella* associated with persistence (23, 24) by dual RNA sequencing (RNA-seq). This simultaneous analysis of the transcriptomes of the host and the pathogen revealed that this specific bacterial subpopulation secrete effectors implicated in reprograming macrophages to promote anti-inflammatory macrophage polarization linked with bacteria long-term survival (25). While the sensitivity of transcriptome analysis outperforms proteome studies, especially in the context of infection, the lower growth rates typically associated with infection (26), bacterial virulence (27, 28) and persistence, are associated with a considerably higher gene expression noise (29) and extensive post-transcriptional regulatory control. Therefore, mRNA levels in bacterial populations with reduced growth rates often correlate poorly with protein levels, stressing the need for protein-based studies. However, as the theoretical *Salmonella* proteome (^~^4,500 proteins) is vastly underrepresented in the context of the infected host (^~^20,000 proteins), especially when considering an up to ^~^1500 fold difference in the *Salmonella* versus human cellular total proteome content in the context of infection (30), studying both proteomes simultaneously and interactively entails proven technical challenges (30, 31), further complicated by the considerable heterogeneity in bacterial and host subpopulations as mentioned above.

By significantly improving peptide detection and quantification, we recently reported on the hybrid use of data-dependent and data-independent acquisition spectral libraries to empower what we called dual proteome profiling (31). In our current study, we used this optimized workflow to perform time-dependent dual proteome profiling of infection-relevant *S*. Typhimurium pathogen encounters with its host, without the need of *a priori* bacterial pathogen enrichment. More specifically, through the use of an intracellular epithelial infection model (i.e., *S*. Typhimurium infected HeLa cells), dual proteome profiling unveiled for the first time a detailed and integrative proteomic perspective on host-pathogen interactions over the time course of an infection. *S*. Typhimurium invasion and replication specific proteomic signatures in epithelial cells revealed interdependent host/pathogen specific responses and pointed to putative novel infection markers and signalling responses.

## Results

### Epithelial Salmonella infection model

In line with previous reports in HeLa cells, while at lower multiplicities of infections (MOI or ratio of bacteria to cells of 10) only ^~^5% of HeLa cells were infected, and since at modest MOI increases (up to MOI 60) higher bacterial loads rather than an increase in the overall number of infected cells has been reported due to *Salmonella* cooperative entry, an MOI of 100 resulted in the infection of over half of the cell population (data not shown) (18, 32–34) and was opted for dual proteome profiling (see below results). Bacterial colony forming unit (CFU) enumeration indicated a ^~^7-fold increase of intracellular bacteria when comparing early versus later times post infection (2 and 8 hours post infection (hpi), respectively) (**Supplementary Figure S1A)**. As bacterial load has proven to impact host signalling and host responses (2, 18, 20, 34), we assessed host cell viability by means of MTT (3-(4,5-dimethylthiazol-2-yl)-2,5-diphenyltetrazolium bromide) and LDH (lactate dehydrogenase) assays (**Supplementary Figure S1B** and **S1C**), demonstrating that the infection conditions used did not significantly affect host cell viability up till 8 hpi when compared to non-infected controls and HeLa cells infected with a non-flagellated, non-invasive *ΔprgH Salmonella* control strain (35).

### Dual proteome profiling of Salmonella-infected epithelial cells

Proteome samples of *S*. Typhimurium infected human HeLa cells were prepared to monitor differences in steady-state protein expression levels at 2, 4, 8, 16 and 24 hpi in biological quadruplicates. Following trypsin digestion, each sample was analysed by LC-MS/MS in both DDA and DIA mode and data analysed respectively using MaxQuant (36) against a composite database containing *Salmonella* and human UniProtKB protein entries, and EncyclopeDIA (37) using our optimized hybrid spectral library workflow (31). The latter combines experimental libraries with predicted spectral libraries to optimize the search space for identifications, shown to facilitate dual proteome profiling (31) and suited for DIA-based exploration of *S*. Typhimurium infected epithelial host cells.

When considering all samples analysed in both DDA and DIA acquisition modes, a total of 7,821 proteins were identified, comprising 6,830 human proteins (87% of all protein identifications) and 991 *S*. Typhimurium (13% of all protein identifications and 21% of the annotated *S*. Typhimurium proteome) proteins.

In a bit more detail, by DDA, 6,696 unique proteins (i.e., 767 *S*. Typhimurium and 5,929 human proteins) (**Supplementary Table S1A**) and, by DIA, 6,734 unique DIA proteins (i.e., 853 *S*. Typhimurium and 5,881 human proteins) (**Supplementary Table S1B**) were identified, implying that the number of human proteins identified remained by and large unaffected when comparing DDA versus DIA data, while a notable increase in *S*. Typhimurium proteins identified (^~^10%) was observed (**Fig. 1A**). The latter is further strengthened when considering peptide to spectrum matches (PSMs) and unique number of peptides identified (**Supplementary Tables S1A-B**).

**Figure 1.**
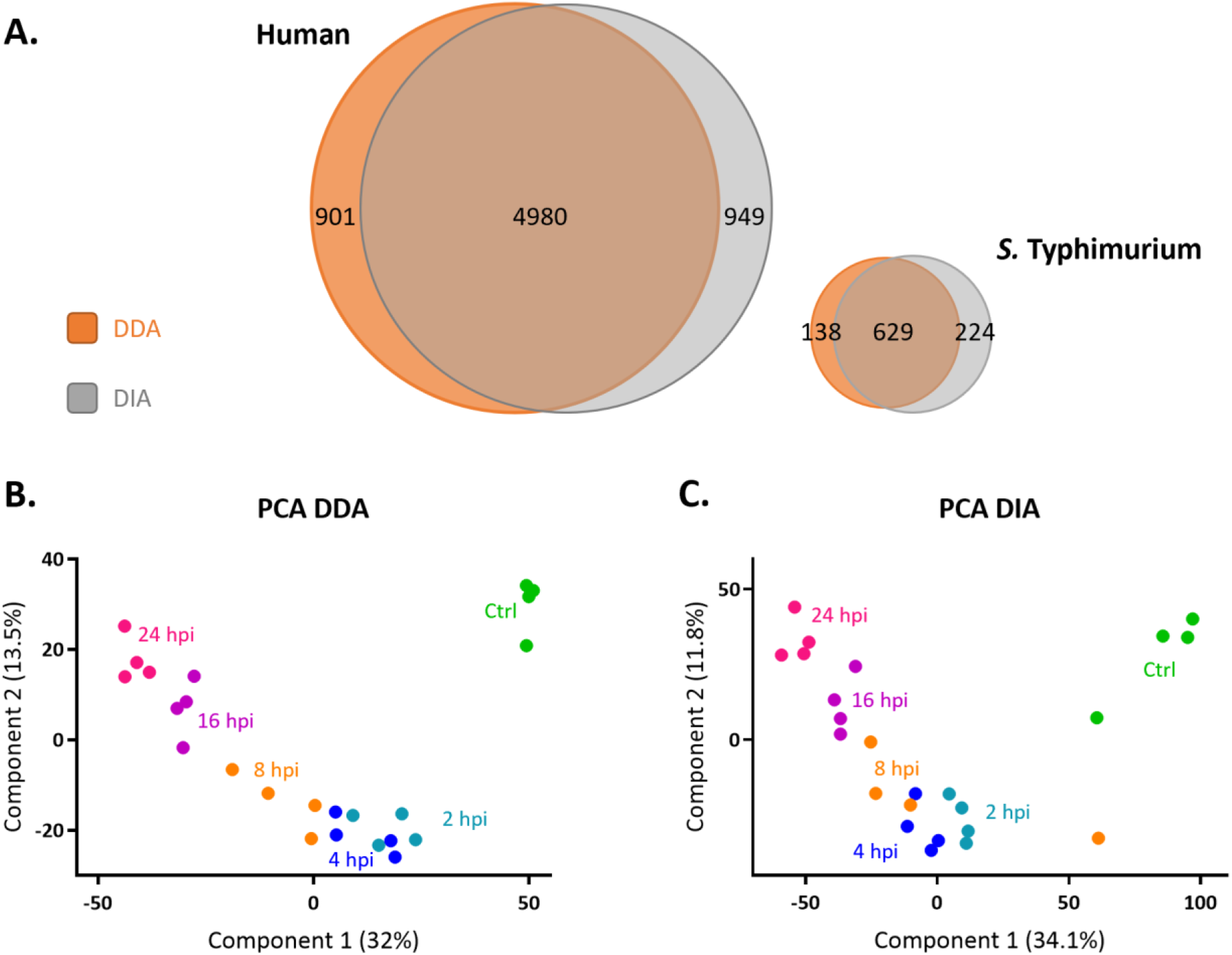
DDA and DIA dual proteome profiling of *Salmonella-infected* epithelial cells. **(A)** Venn diagram created using Venny (http://bioinfogp.cnb.csic.es/tools/venny/index.html) showing the total number of human (left Venn diagram) and *S*. Typhimurium (right) proteins identified using DDA (orange) and DIA (grey) data (both at FDR ≤ 0.01). Variability of DDA **(B)** and DIA **(C)** replicate samples represented in PCA plots. Green, cyan, blue, orange, purple and pink circles represent control (Ctrl), 2, 4, 8, 16 and 24 hpi samples, respectively.

A principal component analysis (PCA) revealed an expected clear variability between control versus *S*. Typhimurium infected HeLa cells as well as a temporal response. Besides clustering according to the time-point post infection, a clearer correlation of proteomes from earlier (2, 4 and 8 hpi) versus later time-points post infection (16 and 24 hpi) (**Figs. 1B-C**) was observed.

To monitor differences in protein levels between all DIA and DDA setups and replicates analysed, multiscatter plots of protein intensity values (log2) were generated. When grouping replicate samples per setup, the average intra-setup Pearson correlations appeared higher for DDA versus DIA (average and minimum correlation of 97% for DDA samples, while average and minimum correlations of 95% and 92%, respectively for DIA samples), and correlation ranged between 92% and 98% for all DDA data, and between 84% and 97% for the DIA datasets when considering average correlations among all different setups (inter-setup) analysed (see correlation plots in **Supplementary Figures S2–S3**).

Further, considering the protein expression ranges observed, a dynamic range spanning over 5 orders of magnitude (max. ^~^365,000-fold change) for DIA data, while only over 4 orders for DDA data was detected (max. ^~^57,000-fold change) (**Supplementary Tables S1A-B**).

### Improved sensitivity of DIA to study Salmonella protein downregulation during infection

After filtering for valid values (minimum 3 valid values in at least one group/setup), median normalized expression per species and imputation of missing values, human and *S*. Typhimurium protein expression values were compared in a pairwise manner with the HeLa control and the *S*. Typhimurium proteomes at 2 hpi, respectively. This selection left 4,666 human (**Supplementary Table S2A**) and 479 *S*. Typhimurium (**Supplementary Table S2B**) protein groups for comparison in case of the DDA data, and 5,818 human (**Supplementary Table S2C**) and 844 *S*. Typhimurium **(Supplementary Table S2D**) protein groups for comparison in case of the DIA data. In line with our dual proteome profiling results reported using artificial proteome mixtures (31), and while the *S*. Typhimurium protein identification rates in DIA only modestly increased (767 vs. 853 *S*. Typhimurium protein identifications), the number of reliably quantified *S*. Typhimurium proteins drastically increased with 76% (1.8-fold) when using DIA data (479 vs. 844 quantified proteins). Further, from the differential expression analysis, it is clear from the corresponding profile plots that both up-as well down-regulation (compared to 2 hpi) of *S*. Typhimurium proteins can comprehensively be studied with DIA data, while expression analysis is mostly confined to upregulated *S*. Typhimurium proteins with DDA data (**Figure 2**). This discrepancy can be explained by the limited DDA over DIA sensitivity and the low *S*. Typhimurium protein content in the proteome samples of infected cells, especially at the earliest time points post infection.

**Figure 2.**
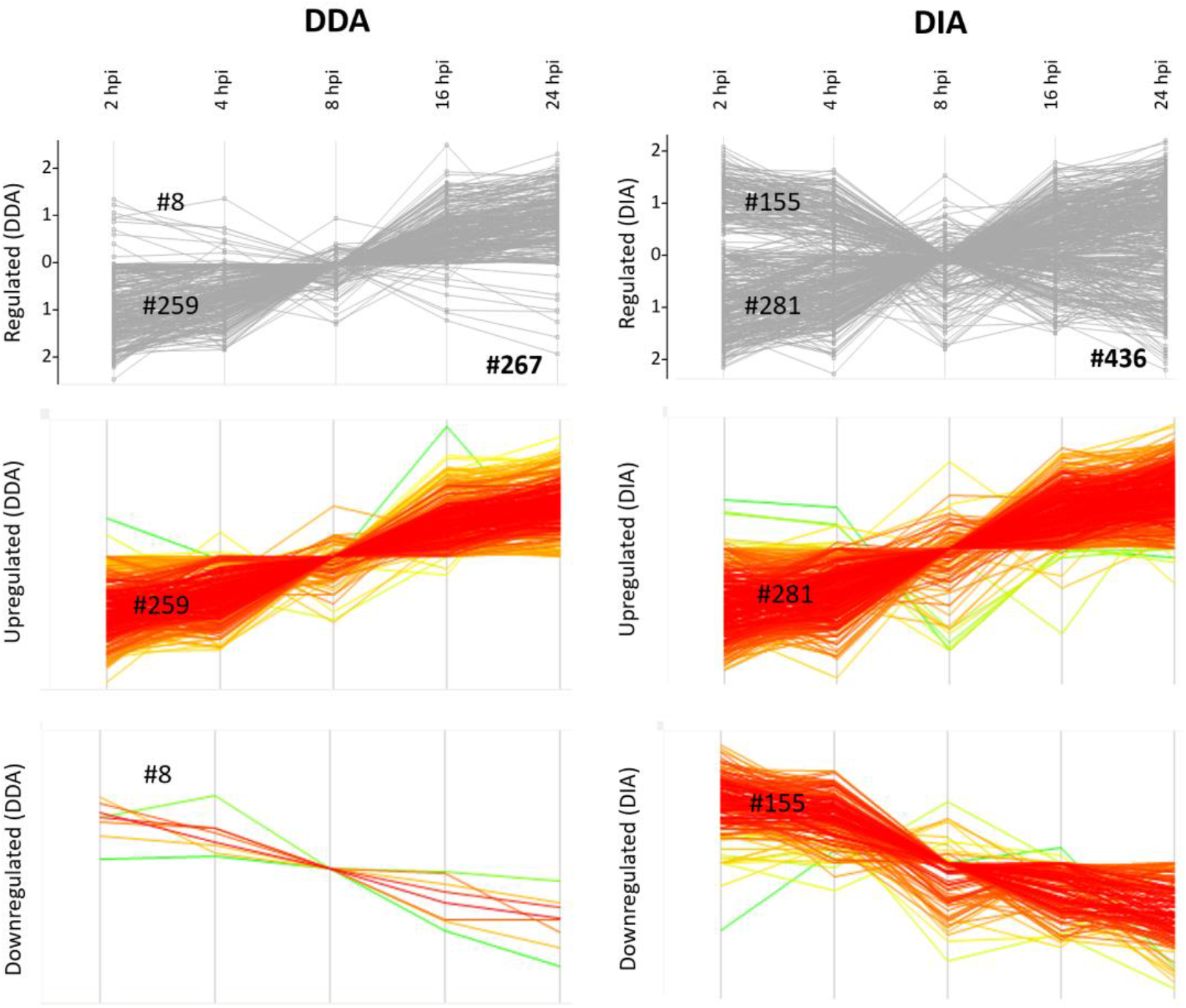
Profile plots of regulated *S*. Typhimurium proteins over the time-course of infection. Significant regulation (FDR ≤ 0.01) was determined across the 5 timepoints post-infection using multiple t-testing and corresponding normalized averaged z-scores of *S*. Typhimurium DDA (left panels) and DIA data (right panels) plotted, with corresponding members of the two main clusters (i.e., up- and downregulated cluster) shown and its corresponding number of regulated *S*. Typhimurium protein groups indicated.

Consequently, studying DDA-based *S*. Typhimurium protein downregulation during infection is rather limited to only a few (8) highly expressed *S*. Typhimurium proteins already identified and quantified at 2 hpi. Because of this finding, we further focussed our dual proteome expression analysis to DIA data. Nonetheless, generally similar trends and overlapping regulations could be observed among the DDA and DIA data (**Supplementary Tables S1, S2A-B** and data not shown). The intensities of significantly regulated proteins after multiple sample ANOVA testing (FDR ≤ 0.01 and an S0 of 0.1) are shown as heat maps and/or profile plots for *S*. Typhimurium and human regulated proteins, respectively (**Figures 2** and **3,** and **Supplementary Tables S2C** and **S2D**). Overall, 1,742 human and 436 *S*. Typhimurium protein groups showed a significantly regulated temporal response in DIA data during the time-course of infection (FDR ≤ 0.01).

**Figure 3.**
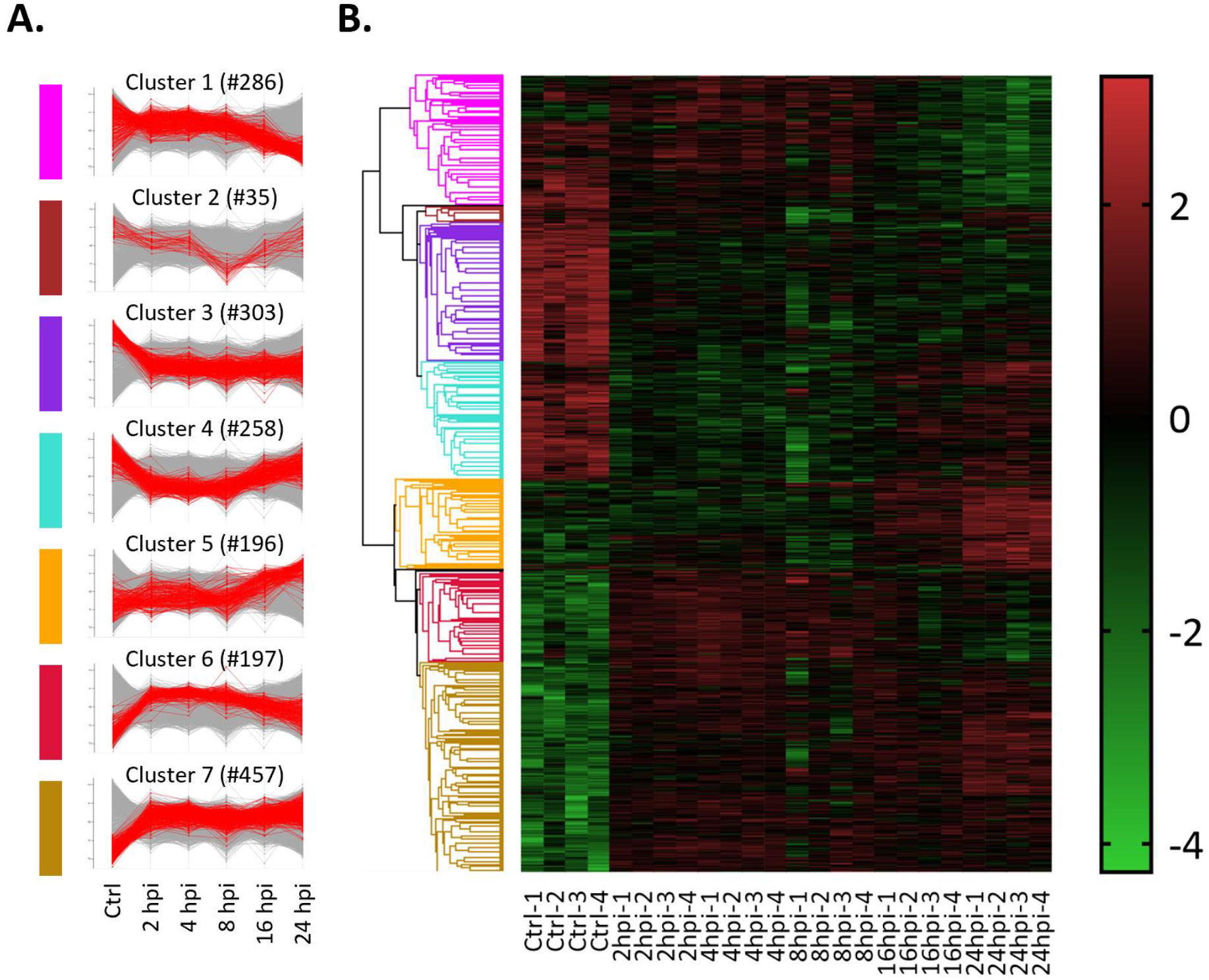
Profile plots and heat map representation of regulated human proteins over the time-course of infection. Regulation was determined across the 5 timepoints post-infection and control cells using t-testing for normalized DIA data. The intensities of significantly (FDR ≤ 0.01) regulated human proteins (**#1,742**) from the DIA analysis are shown. The averaged expression values of corresponding members of the 7 main clusters after ANOVA are shown as profile plots **(A)** and as a heat map **(B)**, with the corresponding number of regulated protein groups indicated per cluster (see also **Supplementary Table S2C**). In panel **B**, green indicates low intensities while red indicates high intensities.

### Host cell responses to Salmonella infection reveal induction of host immunity and epithelial cell differentiation

To map the epithelial host cell response to *S*. Typhimurium infection over time, a gene ontology enrichment analysis (GOBP (Biological Process), GOCC (Cellular Component), GOMF (Molecular Function), general GO and UniProt keywords with expression changes (FDR 0.02)) of the significantly regulated proteins identified in the DIA-MS analysis was performed (**Figure 4** and **Supplementary Table S3)**.

**Figure 4.**
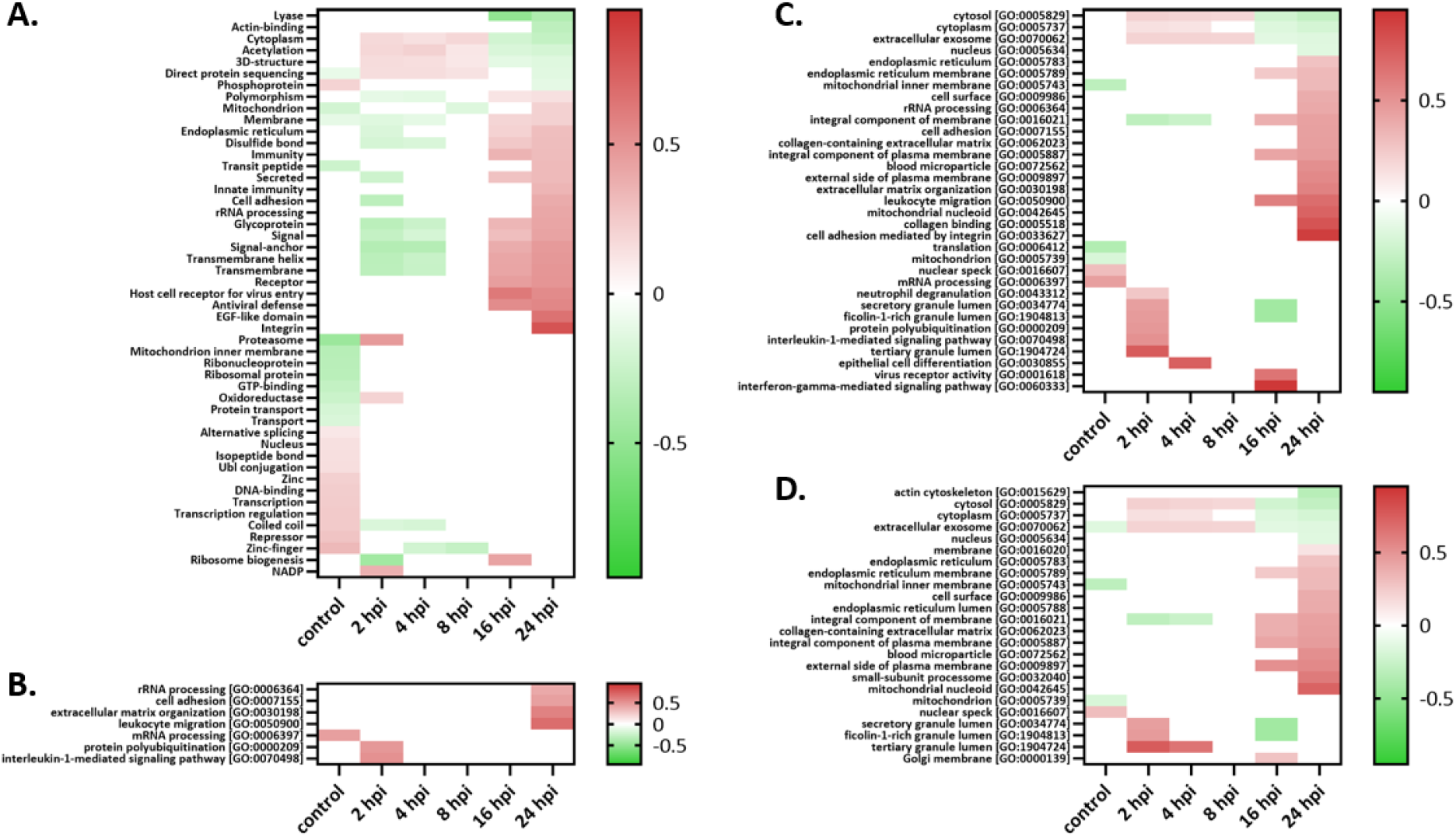
Heatmaps of significantly regulated (FDR ≤ 0.02) normalized human protein annotation enrichment scores across all six conditions (averaged expression values) assayed by data-independent acquisition (DIA). Term enrichment was determined using the 1D annotation enrichment algorithm embedded in the Perseus software suite and p-values were corrected for multiple hypotheses testing using the Benjamini and Hochberg false discovery rate. Enriched annotation terms (FDR ≤ 0.02) in the categories **(A)** UniProt keywords, **(B)** GOBP (Biological Process), **(C)** general GO and **(D)** GOCC (Cellular Component) are shown as heatmaps and colour coded according to the 1D enrichment scores calculated (38), with red and green colouring indicating enriched and depleted annotations, respectively.

At early time points post-infection (i.e., 2 hpi), interleukin-1 mediated signalling and protein polyubiquitination are upregulated indicative of the induction of host immunity, while cell adhesion, extracellular matrix organizations and related terms, e.g., ‘collagen-containing extracellular matrix’ (e.g., including collagen α-3, thrombospondin-1 and Galectin-1) (39) and ‘cell adhesion mediated by integrin’ (e.g., with up to over 7-fold upregulation of the integrins α-2 (ITGA2), α-5 (ITGA5), α-V and β, besides ICAM-1, ADAM9 and vitronectin)) represent upregulated processes opposed to the intracellular actin cytoskeleton downregulation observed at late time points post infection. Some selected representative protein profile plots of the aforementioned significantly regulated categories are shown in **Supplementary Figure S4.** Jointly, these cellular and phenotypic hallmarks reflect epithelial cellular differentiation induced early upon infection (40). The marked induction of host immunity, host integrin signalling and extracellular matrix organization is also in line with previous proteomic reports of infected epithelial cells (41). Overall, besides demonstrating a specific and complex response to bacterial infection over time, our findings seem to indicate that cell differentiation may generate physiological heterogeneity in *S*. Typhimurium infected epithelial cells as previously shown for the *S*. Typhimurium mediated reprogramming of macrophage hosts towards alternative anti-inflammatory (i.e., M2) polarized macrophages, a pathogenic strategy enabling (a subpopulation of) intracellular bacteria to survive and reprogram their host cell to enable bacterial survival and pathogenic spread (25).

### Temporal Salmonella proteome adaptations to adjust to the cytosolic lifestyle in infected epithelial cells

Reflecting the virulence lifestyle of infecting *Salmonella*, the pronounced activity of the two-component system *phoPQ* known to be exerted at later time-points post infection was evidenced by the upregulation of its two components PhoP and PhoQ at later time points post-infection and its downstream impact on SPI-1 and SPI-2 expression more generally (42). More specifically, an enrichment of *ΔphoPQ* regulon targets (43) (i.e. a group of genes regulated as a unit, here genes regulated upon deletion of *phoPQ*) found upregulated (i.e, SipC, SopB, SipB, InvB, PrgH, InvG, PrgI) or downregulated (SodCa, PagC, PhoN, WrbA, UgpB, PdgL, AroQ, PotD, Rna, MsrA, YdcW, SseJ, Tal, DkgA, YdcR, PhnO, SseL) was respectively observed at early (2 hpi) and late (16-24 hpi) times post-infection (FDR ≤ 0.05) (**Supplementary Table S2D** and **Figure 5**). These time-dependent PhoPQ dependencies are in line with the well-established positive regulation of PhoPQ on SPI-2 expression. Other temporal *Salmonella* proteome adaptations were indicative of other affected regulons, with an overall positive regulation of SPI-1 (e.g. *hilA, hilC, fliZ*) or SPI-2 (e.g. *ssrA, slyA*) at early and later time points of the infection, respectively (**Figure 5**).

**Figure 5.**
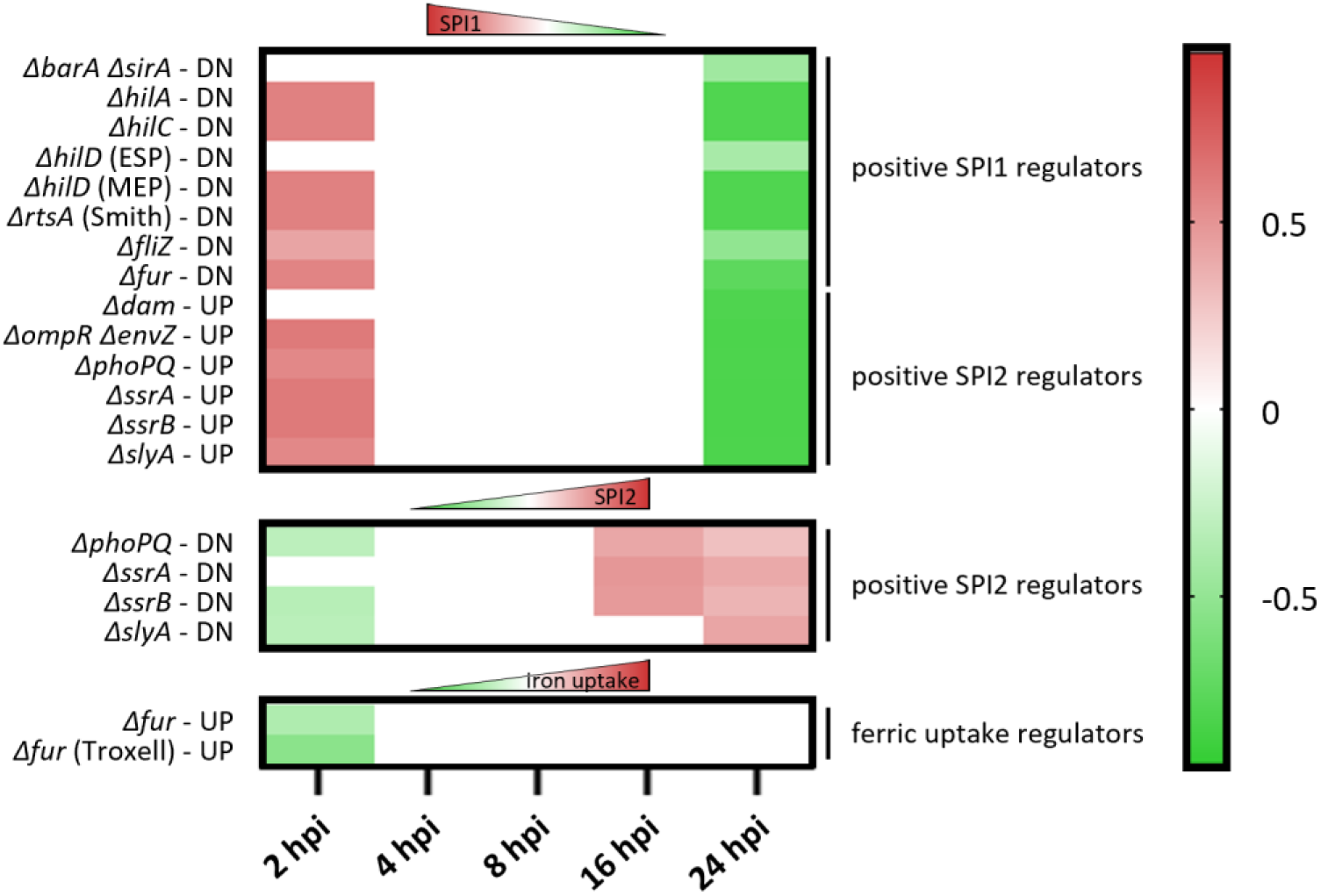
Heatmaps of significantly regulated (FDR ≤ 0.05) normalized *S*. Typhimurium regulon enrichment scores across all infection conditions (averaged expression values) assayed by data-independent acquisition (DIA). Using a recently reported compilation of highly curated regulon gene targets identified by means of microarray or NGS-based gene expression studies (43), enrichment of specific regulons was determined using the 1D annotation enrichment algorithm embedded in the Perseus software suite. P-values were corrected for multiple hypotheses testing using the Benjamini and Hochberg false discovery rate. Enriched annotation terms (FDR ≤ 0.05) in the categories are shown as heatmaps and colour coded according to the 1D enrichment scores calculated (38), with red and green colouring indicating enriched and depleted annotations, respectively. ESP (early stationary growth phase), MEP (mid exponential phase), upregulated (UP) and downregulated (DN) genes as reported in (43).

Further, indicative of the metal ion shortage encountered by the bacterial pathogen when residing in the host (43, 44), significant upregulation of metal ion transporters and uptake systems for zinc (ZnuA), iron (IroBN, EntBF, SitAB, Fur, Mrp, FepAB, CirA) and manganese (SitAB) was observed (**Supplementary Table S2D**). Also, expression of the ferric uptake regulator Fur, which binds to promoters of target genes implicated in iron acquisition and transport in Fe^2+^-rich conditions thereby preventing target transcription, was lowered. In accordance, a significant impact on the *fur* regulon (including *ugpB, sufA, aldB, ynhA* and *tal* as well as the aforementioned *sitAB, cirA, entF, fepA, iroB, entB* targets) was detected, as their corresponding target protein products were found enriched at 24 hpi (FDR ≤ 0.01) (**Figure 5** and **Supplementary Table S2D**). In line with the studies by Liu *et al*. (45) and Li *et al*. (46), bacterial proteome adaptations revealed downregulation of chemotaxis (e.g., CheAMW) and upregulation of His biosynthesis (HisABCDFGHI). In agreement with the increased proliferation rate of cytosolic *Salmonella*, ribosomal proteins (rplBEKMU and rpsCDEJLMRST) and proteins implicated in cell division (MinE, MpI, MukB, MurAEFG, Pal, TolB) were found upregulated at later timepoints of the infection (**Supplementary Table S2D** and **Supplementary Figure S5**).

Since extensive transcriptional reprogramming was shown to accompany the successful colonization of the epithelial cytosol, a niche occupied by a hyper-replicating *Salmonella* sub-population especially at later times post-infection, it is noteworthy that upregulation of cytosol signature genes previously reported to be transcriptionally upregulated in Gram-negative pathogens colonizing the cytosol were also observed as being regulated in our proteome study (i.e., *bioA, entF, fepA, fepB, iroB, iroN* and *sitAB*). Furthermore, Powers *et al*. (43) demonstrated that a *S*. Typhimurium *ΔsitAΔmntH* mutant was compromised for growth in the cytosol of epithelial cells linking this to the need for acquiring Mn^2+^ acquisition when residing in this subcellular location. Altogether, proteome profiling of infecting salmonellae is heavily biased towards its virulence repertoire and provides a proteomic perspective on host-pathogen interactions and pathogen localisation over the time course of infection.

## Discussion

Given the generally acknowledged multifactorial complexity of host-pathogen interactions, the variability in the results from infection studies reported, and the high temporal aspect of the interactions, multiplex omics-based analyses deliver a comprehensive understanding of the complexity of such systems and the interplay between pathogen and host. To understand bacterial adaptation to host encounters, to date, the vast majority of *Salmonella* centred proteomics research relied on axenic grown *Salmonella* exposed to various (infection-relevant) growth conditions (47). Only more recently, extensive adaptation of *S*. Typhimurium to infected host epithelial cells was shown by proteomics efforts making use of enriched *Salmonella* isolated from infected epithelial host cells (44–46). Here, we report for the first time a dual proteome profile of *Salmonella*-infected epithelial host cells in function of time, both in DIA and DDA modes, overall providing a proteomic perspective on the bacterial pathogen encounter with its epithelial host and permitting the simultaneous study of host and pathogen proteomes.

Noteworthy, while DIA outperformed DDA both in terms of identifications and quantifications, especially for the bacterial pathogen, for DDA data, correlation of quantifications was shown to be higher than for DIA data. This finding is mainly attributed to the increased DIA-based quantification of low abundant peptides and due to the higher variability of quantification data for low abundance species given the interference with random noise (31).

This observation is also apparent from the generally less discrete (differential) patterns of *Salmonella* protein expression as compared to the distinctive human expression clusters. The former finding is also likely (in part) due to the variability of proteome sampling at different timepoints post-infection. Bacterial proteome sampling in dual proteome profiling nevertheless remains less comprehensive as compared to previously reported protein identifications making use of pure *Salmonella* cultures or of *Salmonella* isolated from infected epithelial host cells. For example, when considering axenic *S*. Typhimurium cultures and triplicate shotgun proteome analysis, typically a 2-fold increase in the number of *S*. Typhimurium proteins identified and quantified (48) (around ^~^1,500 to 1,600) are found using similar MS instrumentation in DDA mode (30).

However, the high quantification sensitivity obtained by DIA permitted the reliable capture of over 100 regulated bacterial proteins over the time course of infection, generally exceeding the numbers of regulated genes reported in other studies, and, importantly, here also including the identification of downregulated *Salmonella* proteins upon infection.

By comparing protein expression regulation across infection conditions only, and while initial bacterial adaptations to the host might be missed, upregulation of e.g., phosphate utilization, previously shown to be highly induced in the *Salmonella* proteome upon medium switching occurring prior to bacterial internalization (44), was not observed in our study. This indicates that likely phosphate shortage is not experienced within the host itself but rather dependent on bacterial cultivation steps preceding the actual infection, necessitating the need for detailed experimental descriptions, frequently lacking in literature, and the need for uniformity when comparing omics datasets, hence the need for dual omics strategies.

On a related note, upon systematically comparing our dual proteomics data with previously reported transcriptome changes making use of dual RNA-sequencing and FACS-enriched *S*. Typhimurium infected HeLa cells (49), for the about 5,700 matching identifications (DIA data), transcriptome versus proteome changes were found to only poorly correlate, especially at later timepoints post-infection (Pearson correlation below 0.2 at 24 hpi). Annotation enrichment analysis even pointed to oppositely regulated gene expressions when comparing protein versus transcript levels (data not shown). In particular, while proteome data indicates downregulation of the GO term ‘actin cytoskeleton’ at 24 hpi, upregulation was observed at the transcript level. Opposingly at 16 and 24 hpi, upregulated ribosome biogenesis and mitochondrial gene expression was observed at the proteome level, while transcript levels were down, while at earlier time points post-infection, a reverse proteome versus transcriptome regulation was observed for ribosomal proteins being down and transcripts up. Overall, this indicates that besides the lag phase between transcription and translation to be considered, translational and/or post-translational control significantly impacts bacterial gene expression, especially in the context of (prolonged) infections, further emphasizing the need to study gene expression changes at the proteome level.

Further noteworthy in this context is that *S*. Typhimurium T3E host target proteins were found specifically enriched in the category of regulated proteins as determined by a chi-square test of independency (p < 0.01; p-value 0.002203), indicating that bacterial effector functioning greatly impacts the proteome of the host. As a representative example, it has been shown that *Salmonella* remodels the host actin cytoskeleton to force its entry into non-phagocytic cells. These cytoskeletal rearrangements are mediated by T3SS-1 effectors, such as SopB, SopE and SopE2 with indirect actions that are essential for efficient invasion (50). Several other effectors were shown to directly interact with actin(-binding) proteins including effector SspH2 that has established interactions with profilin (PFN)-1 and −2 inside the host (51–53). In our DIA data, PFN2 showed to be ^~^3-fold upregulated at 2 hpi. With its predicted inhibitory effect on the formation of membrane ruffles, PFN2 upregulation at 2 hpi supports the predicted downregulation of “formation of cellular protrusions” when performing Ingenuity Pathway Analysis (IPA) (**Figure 6A**).

**Figure 6.**
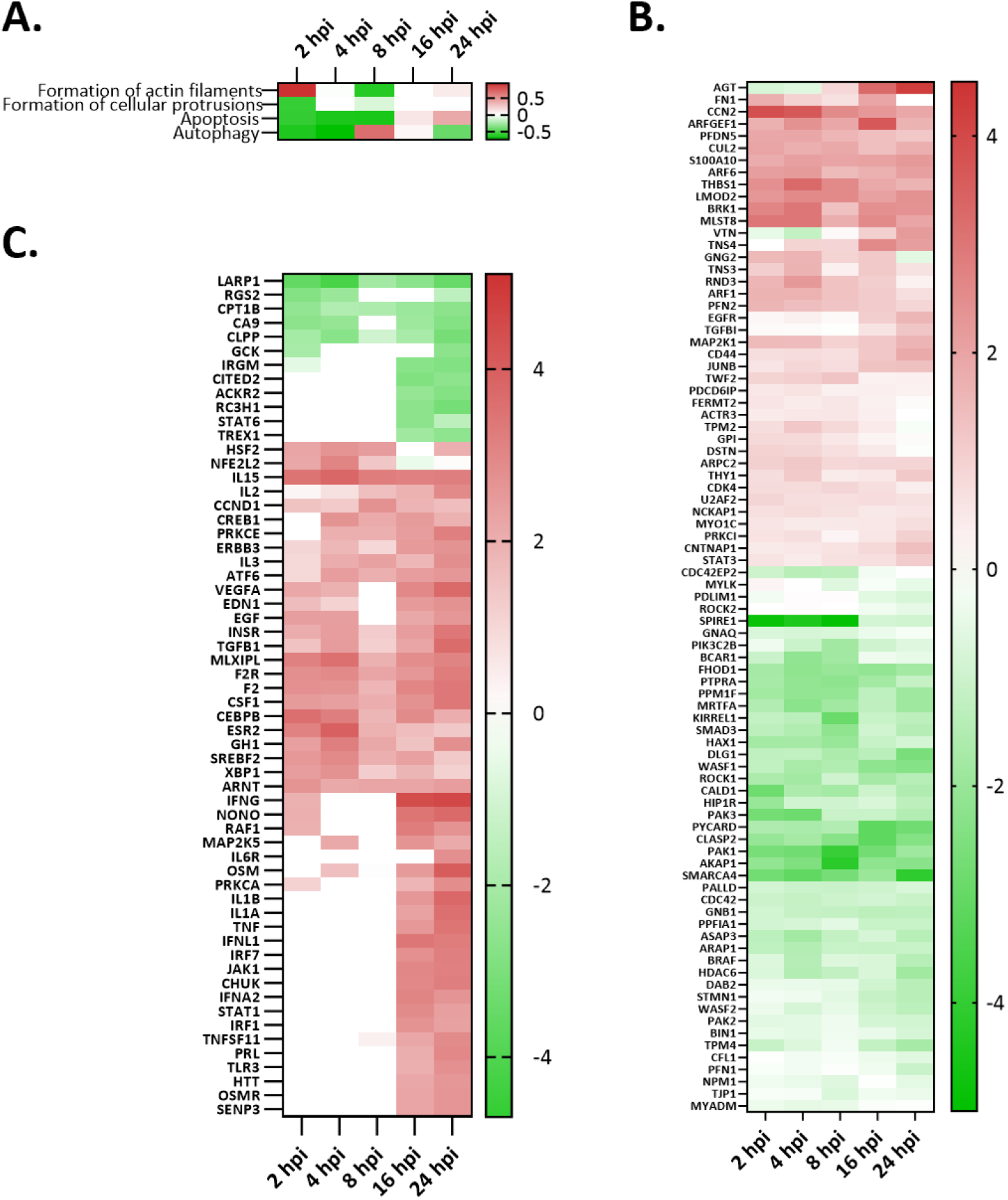
Selection of functions and regulators predicted to be activated or inhibited using Ingenuity Pathway Analysis (IPA). **(A)** Heatmap visualization (IPA activation z-score) of selected IPA functions regulated over the course of the infection. **(B)** Overview of regulated genes (#85) (log FC) contributing to the IPA function “formation of actin filaments” (see also **Supplementary Table S2C**). **(C)** Overview of activated and inhibited upstream regulators. Heatmap representation of 60 predicted proteinaceous (activated or inhibited for at least one of the timepoints post-infection analysed) upstream regulators (p-value ≤ 0.01), colour coded according to the IPA activation z-score.

To restore the host cell cytoskeleton, *Salmonella* effector SptP uses its GTPase-activating protein (GAP) activity to inactivate RAC1 (54), thereby reversing the Rho GTPase activating effects of SopE (55). This restoration of the host cytoskeleton is also evident from the predicted activation of the “formation of actin filaments” at 2 hpi and the regulation of contributing genes (**Figure 6A-B**).

Further, IPA pathway analysis substantiated the observation that apoptosis of epithelial cells infected with *Salmonella* is delayed for until 8 hpi (**Figure 6A**) (56), an observation in line with our host cell viability data. This delay in apoptosis is thought to be attributed to sustained AKT signalling that is accredited to multifaceted effector SopB, a lipid phosphotransferase (57).

Further, in order for *Salmonella* to manipulate the host endocytic pathway, several ras-associated binding (RAB) proteins have been reported to be directly targeted by effectors (e.g., GtgE, SopD and SopD2 (58–60). Based on our IPA analysis, autophagy has a positive activation z-score at 8 hpi (**Figure 6A**) with a predicted contribution of the RAB1A, RAB7A and RAB9A found to at higher abundancies at this time point in our DIA data.

Next to demonstrating the fluctuations among functions affected during *Salmonella* infection, IPA revealed 60 putative upstream regulators to be either activated or inhibited over the course of infection (**Figure 6C**). Among others, immune system signalling regulators such as toll-like receptor signalling proteins, e.g. STAT1, IL1B, IFNA2 and TLR3 and MAPK3, were shown to be activated at 16-24 hpi.

With some of the expression differences linked to intracellular host localization of the bacteria and shown to promote intracellular *Salmonella* growth (45), studies on the causal links between host/pathogen expression interdependencies remain an interesting area of future research.

With the observed variability of *Salmonella* subpopulations observed in diverse host cells (23, 24), future DIA-based dual proteome profiling infection studies performed at the sub-population or even single-cell level may further advance our understanding of host-pathogen interactions at an unprecedented resolution.

## Materials and Methods

### Bacterial strain and growth cultivation conditions

The *Salmonella enterica* serovar Typhimurium (*S*. Typhimurium) wild-type (WT) strain SL1344 (Hoiseth and Stocker, 1981) (Genotype: hisG46, Phenotype: His(-); biotype 26i) was obtained from the *Salmonella* Genetic Stock Center (*Salmonella* Genetic Stock Center (SGSC), Calgary, Canada; cat n° 438 (61)). Bacterial growth was performed in Luria Beltrami (LB)-Miller broth (10 g/L Bacto tryptone, 5 g/L Bacto yeast extract, 10 g/L NaCl). For bacterial cultivation, single colonies were picked from LB plates, inoculated in 3 ml liquid Lennox (L) growth medium (L-broth) in round-bottom falcon tubes (Corning, cat n° 352059) and grown for ^~^18 h at 37 °C with agitation (180 rpm). Subsequently, the cultures (^~^OD_600_ 2.0) were diluted 1:50 in T25 flasks with ventilated cap in 8 ml L-medium and grown at 37 °C and 180 rpm to early stationary phase (OD_600_ 3.0) (corresponding to ^~^2 x 10^9^ bacteria per ml and ^~^3.5 h of culture after 1/50 dilution of an overnight culture).

### Cell culture

Human epithelial HeLa cells (epithelial cervix adenocarcinoma cell line, American Type Culture Collection, Manassas, VA, USA; ATCC^®^ *CCL-2*™) were cultured in GlutaMAX containing Dulbecco’s Modified Eagle Medium (DMEM) (Gibco, cat n° 31966-047) supplemented with 10% foetal bovine serum (FBS) (Gibco, cat n° 10270-106) and 50 units/ml penicillin and 50 μg/ml streptomycin (Gibco; cat n° 5070-063). Cells were cultured at 37 °C in a 5% CO_2_ gas-equilibrated humidified incubator and passaged every 3-4 days.

For infection, HeLa cells were seeded in a T175 (Greiner Bio One, cat n° 660160) at a density of 10^7^ cells in 19 ml of medium for dual-proteomics or at 1.5 x 10^4^ cells per well in 96-well plates (Nunc, cat n° 164590) in 0.2 ml of medium for MTT, LDH assays and CFU enumeration. The culture medium was supplemented with 10% heat-inactivated FBS (heat inactivation was done for 30 min at 56°C) and without antibiotics. In both set-ups, HeLa cells were seeded one day prior to infection thereby reaching a final confluence of ^~^85-90%, corresponding to approx. 1.25 x 10^7^ HeLa cells per T175 cell culture flask and 1,875 x 10^4^ HeLa cells per well in 96-well plates at the day of infection.

### Bacterial infection of cultured HeLa cells

For bacterial invasion, the bacteria were harvested for infection at early stationary phase (when studying *Salmonella* spp. invasion of epithelial cell lines, the early stationary phase was shown to be the phase during which *Salmonella* pathogenesis island-1 (SPI-1) is highly induced, and therefore a suitable growth phase for infection (62)) after reaching an optical density at 600 nm of 3.0. Bacterial cells were collected by centrifugation (6,000 × *g*, 10 min) at room temperature, washed once with preheated (37°C) Dulbecco’s Phosphate Buffered Saline (DPBS) (Gibco; cat n° 14190-144), re-spun and resuspended in pre-heated DMEM without serum.

*Salmonella* infection was allowed to proceed in DMEM at a multiplicity of infection (MOI) of 100 (i.e., per T175 cell culture flask of 1,25 x 10^7^ HeLa cells, 1,25 x 10^9^ bacterial cells (or 1.25 ml of a 10^9^ bacteria bacterial cell suspension) were added, or in case of 96-wells, 50 μl containing 1,875 x 10^6^ bacterial cells were added) for 30 min at 37 °C in a 5% CO_2_ gas-equilibrated humidified incubator. After infection, the medium was removed from the culture flasks or the 96-well plates, the monolayers washed with DPBS pre-heated at 37°C and respectively, 20 ml or 0,2 ml of pre-warmed DMEM supplemented with heat-inactivated FBS and 100 μg/ml gentamicin (Sigma-Aldrich, G1264-1G) was added to kill extracellular bacteria after which the infection was allowed to proceed for an additional 2 h. When the infection was allowed to proceed for longer times, after 2 h cells were washed and DMEM supplemented with 10 μg/ml gentamicin was added. For dual proteome profiling, 4 replicate samples corresponding to 2, 4, 8, 16 and 24 hours post-infection (hpi) were collected, besides a HeLa control setup. After a DPBS wash (37 °C) of the cell monolayers, cells were harvested by trypsinization (Trypsin-EDTA; Gibco, cat n° 25300054), collected by centrifugation (1000 × *g*, 5 min) at 4 °C and the cell pellets washed 2 times with DPBS, flash frozen in liquid nitrogen and kept at −80 °C until further processing.

### MTT and LDH assays

For MTT assays, 4 replicates samples were analysed at 2 and 8 hpi. At each time point, culture media was replaced by culture media supplemented with 1 mg/mL MTT reagent (Sigma, cat n° M5655). MTT reduction to formazan crystals by cellular oxidoreductases were allowed to form by incubating the cells in a 5% CO_2_ gas-equilibrated humidified incubator at 37°C for 2 h. Subsequently, media was removed and crystals dissolved in 1 mM isopropanol 1N HCl solution for 30 min when shaken at 500 rpm. The dissolved crystals were quantified by measuring absorbance at 595 nm reflecting cellular metabolic activity as an indicator of cell viability.

For LDH assays, 8 replicate samples were analysed at 2 and 8 hpi by transferring 50 μL of the culture media into a fresh 96-well plate and LDH enzyme activity (release of LDH enzyme upon cell membrane damage is indicative for cellular integrity and concomitantly cell viability) was measured using the LDH Cytotoxicity Assay Kit (Pierce™, cat n° 88953) following the manufacturer’s instructions.

### *prgH* deletion mutant generation by λ Red recombineering

*S*. Typhimurium SL1344 *prgH* mutant (Δ*prgH*) was constructed by λ Red recombineering (63). Briefly, ^~^2.4 x 10^9^ of λ red-induced (0.2% L - (+) – arabinose w/v) electrocompetent and pKD46 (accession number #7669, Coli Genetic Stock Center (CGSS), Yale University, USA) transformed *S*. Typhimurium WT cells (culture grown to OD_600_ 0.6) were electroporated with a linear PCR-editing substrate designed to replace *prgH* with the kanamycin-resistance cassette from pKD4 (accession number #7632, CGSS, Yale, USA). PCR was used to amplify the antibiotic-resistance cassette with 5’ and 3’ 50 bp homology arms complementary to the flanking regions of *prgH* using primer sequences caatggggatgatggttcttttaatatgtgttgagacgcattatacagaat*gtgtaggctggagctgcttc* and acgccgccagtagcgccggatcggagggttttgctgctaatttatccagctatgaatatcctccttagttcctatt). Immediately following electroporation, bacterial cells were recovered in pre-warmed (28°C) Super Optimal broth with Catabolite repression (SOC, consisting of 2% tryptone, 0.5% yeast extract, 10 mM NaCl, 2.5 mM KCl, 5 mM MgCl2, 10 mM MgSO4 and 20 mM glucose) media at 28°C and incubated at 180 rpm for 1 h and subsequently at 37°C for 3 h. Mutant colonies were selected after plating the cell suspension and overnight incubation (upside down) on LB agar plates supplemented with 25 μg/ml kanamycin at 37°C. Allelic replacement of *prgH* was confirmed by colony PCR using primer sequences ggatgatggttcttttaatatgt and ggcaagaaagccatccagtttactttgca, and acgccgccagtagcgccggatcg and gacgagttcttctgagcgggact, and the resulting PCR products sequence verified by Sanger sequencing (Applied Biosystems 3730xl, Thermo, VIB Genomic Service Facility, University of Antwerp, Belgium).

### Bacterial enumeration by CFU counts upon host-selective lysis of *Salmonella-infected* HeLa cells

For CFU enumeration, 3 replicate samples were analysed at 2 and 8 hpi. *Salmonella*-infected HeLa cultures (96-well format) washed with 0,2 ml of pre-heated DPBS were added 0,1 ml of ice-cold lysis buffer (1% TX-100 in DPBS) and incubated for 5 min on ice to allow selective host cells lysis. 0,2 ml of DPBS was added to each well, lysates were transferred to Eppendorf tubes and centrifuged for 5 min at 6000 x *g* and 4°C. The supernatant was recovered and 4 serial 5-fold dilutions (1/5 to 1/625) were prepared in DPBS. 100 μL of each dilution was spread on LB-agar plates using glass plating beads (Thermo Fischer Scientific, cat n° MP115000550), incubated upside down overnight at 37°C, and bacterial enumeration performed by viable CFU counting.

### Proteome extractions and sample mixes

Cell pellets of 4 replicate samples and 6 setups (control HeLa + 2, 4, 8, 16 and 24 hpi samples) were resuspended in guanidinium-hydrochloride (Gu.HCl) lysis buffer at 20 x 10^6^ cells/ml (4 M Gu.HCl, 50 mm ammonium bicarbonate (pH 7.9)) and lysed by three rounds of freeze-thaw lysis in liquid nitrogen. The lysates were sonicated (Branson probe sonifier output 40, 50% duty cycle, 3×30 s, 1 s pulses) followed by centrifugation (16,100 x *g*, 10 min at 4 °C) to remove cellular debris. The protein concentration of the supernatant was determined by Bradford measurement (64) according to the manufacturer’s instructions (Bio-Rad, cat n° 5000006).

An aliquot adjusted with lysis buffer to 2 mg/ml, and equivalent of 400 μg of total protein (200 μl) was transferred to a clean Eppendorf tube, twice diluted with HPLC-grade water, and precipitated with 4 volumes of acetone (at-20 °C) overnight. The precipitated protein material was recovered by centrifugation for 15 min at 16,000 x *g* at 4 °C, pellets washed twice with cold 80% acetone, and air dried upside down for ^~^10 min at room temperature or until no residual acetone odour remained. Protein pellets were resuspended in 200 μl TFE (2,2,2-trifluoroethanol) digestion buffer (10% TFE, 100 mM ammonium bicarbonate pH 7.9) with sonication (Branson probe output 20; 1 s pulses) until a homogenous suspension was obtained. All samples were digested overnight at 37 °C using MS-grade trypsin/Lys-C Mix (Promega, Madison, WI) (enzyme/substrate of 1:50 w/w) while mixing (550 rpm). Samples were acidified with TFA to a final concentration of 0.5% and cleared from insoluble particulates by centrifugation for 10 min at 16,000 x *g* at 4 °C and the supernatant transferred to new tubes. Methionine oxidation was performed by addition of hydrogen peroxide to a final concentration of 0.5% for 30 min at 30 °C. Solid phase extraction of peptides was performed using C18 reversed phase sorbent containing 100 μl pipette tips Bond Elut OMIX 100 μl C18 tips (Agilent, Santa Clara, CA, USA, cat n° A57003100K) according to the manufacturer’s instructions. The pipette tip was conditioned by aspirating the maximum pipette tip volume of water:acetonitrile, 50:50 (v/v) and the solvent was discarded. After equilibration of the tip by washing three times with the maximum pipette tip volume in 0.1% TFA in water, 100 μl of the acidified samples (original input of ^~^200 μg) were dispensed and aspirated for 10 cycles for maximum binding efficiency. The tip was washed three times with the maximum pipette tip volume of 0.1% TFA in water:acetonitrile, 98:2 (v/v) and the bound peptides were eluted in LC-MS/MS vials with the maximum pipette tip volume of 0.1% TFA in water:acetonitrile, 30:70 (v/v). The samples were vacuum-dried in a SpeedVac concentrator and re-dissolved in 50 μl of 2 mM tris(2-carboxyethyl)phosphine (TCEP) in 2% acetonitrile spiked with an indexed Retention Time or iRT peptide mix (i.e., a mixture of eleven non-naturally occurring synthetic peptides added according to manufacturer’s instructions (Biognosys)), to enable retention time (RT) prediction (see below). Samples were stored at −20 °C until LC-MS/MS analysis.

### LC-MS/MS data acquisition

For mass spectrometry analyses, 10 μl was injected from each sample for LC-MS/MS analysis on an Ultimate 3000 RSLCnano system in-line connected to a Q Exactive HF Hybrid Quadrupole-Orbitrap BioPharma mass spectrometer (Thermo). Trapping was performed at 10 μL/min for 4 min in loading solvent A (0.1% TFA in water/ACN (98:2, v/v) on a 20 mm trapping column (made in-house, 100 μm internal diameter (I.D.), 5 μm beads, C18 Reprosil-HD, Dr. Maisch, Germany). After flushing from the trapping column, the peptides were loaded and separated on an analytical 200 cm μPAC™ column with C18-endcapped functionality (PharmaFluidics, Belgium) kept at a constant temperature of 50 °C. Peptides were eluted by a non-linear gradient reaching 9% MS solvent B (0.1% FA in water/acetonitrile (2:8, v/v)) in 15 min, 33% MS solvent B in 105 min, 55% MS solvent B in 125 min and 99% MS solvent B in 135 min at a constant flow rate of 300 nL/min, followed by a 5-minutes wash at 99% MS solvent B and re-equilibration with MS solvent A (0.1% FA in water). For both analyses (data-dependent acquisition (DDA) and data-independent acquisition (DIA), a pneu-Nimbus dual column ionization source was used (Phoenix S&T), at a spray voltage of 2.6 kV and a capillary temperature of 275 °C. For the first analysis (DDA mode), the mass spectrometer automatically switched between MS and MS2 acquisition for the 16 most abundant ion peaks per MS spectrum. Full-scan MS spectra (375-1500 *m/z*) were acquired at a precursor resolution of 60,000 at 200 *m/z* in the Orbitrap analyser after accumulation to a target value of 3,000,000. The 16 most intense ions above a threshold value of 13,000 were isolated for higher-energy collisional dissociation (HCD) fragmentation at a normalized collision energy of 28% after filling the trap at a target value of 100,000 for maximum 80 ms injection time using a dynamic exclusion of 12 s. MS2 spectra (200-2,000 *m*/*z*) were acquired at a resolution of 15,000 at 200 *m/z* in the Orbitrap analyser. Another equivalent 10 μL aliquot from each sample was analysed using the same mass spectrometer now operated in DIA mode. Nano LC conditions and gradients were the same as used for DDA. Full-scan MS spectra ranging from 375-1,500 *m/z* with a target value of 5E6 were followed by 30 quadrupole isolations with a precursor isolation width of 10 m/z for HCD fragmentation at a normalized collision energy of 30% after filling the trap at a target value of 3E6 for maximum injection time of 45 ms. MS2 spectra were acquired at a resolution of 15,000 at 200 m/z in the Orbitrap analyser without multiplexing. The isolation intervals ranged from 400 – 900 *m/z* with an overlap of *m*/*z*.

### Data-dependent acquisition (DDA) data processing

For DDA-based peptide identification and quantification, Raw data files were searched by MaxQuant (36) (version 1.6.9.0). The protein search database comprises the UniProtKB *Salmonella* reference proteome concatenated to the human UniProtKB reference proteome (UP000008962 and UP000005640, together 74,449 proteins), as well as MaxQuant built-in contaminants.. In addition, contaminant proteins present in the contaminants.fasta file that comes with MaxQuant and the 11 iRT peptide sequences (Biognosys-11) were added to the search database. Methionine oxidation was set as fixed modification, and N-terminal acetylation was set as variable modification. To augment protein/peptide quantification, we enabled built-in matching-between-run and label-free quantitation (LFQ) algorithms using default settings in MaxQuant. We used the enzymatic rule of trypsin/P with a maximum of 2 missed cleavages. The main search peptide tolerance was set to 4.5 ppm and the ion trap MS/MS match tolerance was set to 0.5 Da. Default PSM, peptide and protein level FDR thresholds of 1% were used, with an additional minimal Andromeda score of 40 for modified peptides. Protein FDR was set at 1% and estimated by using the reversed search sequences. The maximal number of modifications per peptide was set to 5. For protein quantification in the proteinGroups.txt file, razor peptides were considered and further allowed all modifications.

### Data-independent acquisition (DIA) data processing

#### DDA-based spectral library construction

The ‘msms.txt’ files outputted by MaxQuant DDA searches were used as input for creation of redundant BLIB spectral libraries using BiblioSpec (version 2.1) (65). Redundant spectra were subsequently filtered using the ‘BlibFilter’ function, requiring entries to have at least 5 peaks (‘-n 5’). Peptide sequences uniquely identified in only one of the DDA searches were appended to the spectral library to extend the library and to augment the detection capacity of *Salmonella* peptides. We transformed the peptide RTs present in the BLIB library to iRT using the spiked-in iRT peptides (Biognosys-11). To this end, empirical RTs of the top-scoring iRT peptide identifications (lowest posterior error probability, ‘msms.txt’) in the DDA samples, were used to fit a linear trendline and scale RTs. For the artificial mixtures and *Salmonella* pre-fractionation analyses (31), the corresponding trendlines were iRT = 1.220 RT – 74.566 and iRT = 1.189 RT – 75.039. The updated BLIB files were then converted to DLIB format by EncyclopeDIA, using the combined *human-Salmonella* UniProtKB proteome FASTA as background (66).

#### Library-free searching of proteome FASTA by PECAN/Walnut

DIA raw data files were converted to mzML by MSConvert using vendor peakPicking. Pre-processed DIA samples were searched against a compilation of the *Salmonella* UniProtKB proteome (UP000008962, 4,657 proteins) and human Swiss-Prot proteome (UP000005640, 20,367 proteins) using the EncylopeDIA built-in PECAN algorithm (66, 67). We opted to solely search the human Swiss-Prot protein database, which resulted in a ^~^3-fold reduction of the protein database search space (25,024 versus 74,449 proteins when combining *Salmonella* [UP000008962] and human [UP000005640] UniProtKB references proteomes), in order to minimize the theoretical search space. This was desired to limit the size of a predicted spectral library for all possible tryptic peptides and overall runtime and memory usage. Since 36,494 out of 36,668 (i.e., 99.53%) of all human peptides identified by MaxQuant matched a Swiss-Prot protein entry, no drastic loss in identifications is anticipated. Default settings were used, except for methionine oxidation (to methionine-sulfoxide) being set as fixed modification, and considering a maximum length of 25 amino acids and HCD as fragmentation type.

#### Construction of an MS^2^PIP-based spectral library

MS2 spectra were predicted by MS^2^PIP (version 20190312) (68) for tryptic peptides derived from an *in silico* digest of the *Salmonella* UniProtKB proteome and human Swiss-Prot proteome (trypsin/P, peptide length 7-25 AA, mass 500-5,000 Da, one missed cleavage, N-terminal initiator methionine removal considered) in case of 2+ and or 3+ peptide precursor fit within 400 to 900 *m/z* (scanned range DIA). This yields a total of 1,586,777 predicted MS2 spectra for 1,151,386 peptides solely matching human proteins, 197,782 spectra for 144,156 peptides matching *Salmonella* proteins, and 117 spectra for 110 peptides matching both species. We set methionine oxidation (to methionine-sulfoxide) as a fixed modification for MS2 prediction by MS^2^PIP. Predicted spectra were supplemented with DeepLC predicted RTs using a model trained on RTs of 35,206 non-redundant peptides identified in DIA PECAN searches (peptide Q-value < 0.01) (as described in the above section).

#### Hybrid library construction

The chromatogram libraries (ELIB) generated by EncyclopeDIA after PECAN and MS2PIP-library searching were combined into a single redundant library. After conversion to BLIB format, and adding the ‘:redundant:’ tag in library info (sqlite3), the ‘BlibFilter’ function was used to create a non - redundant DIA spectral library as described above. This DIA spectral library was further extended with spectra from the extended DDA spectral library (see above) corresponding to peptides not yet contained within the DIA spectral library. The resulting hybrid DDA/DIA spectral library (i.e., joining DDA- and DIA-based identifications) was then converted to DLIB format as described above.

#### EncyclopeDIA spectral library searching and peptide quantification

The resulting mzML files were searched against the DDA, MS2PIP, or hybrid spectral DLIB libraries using EncylopeDIA software (version 0.90) (66) with default settings. Sample-specific Percolator output files and chromatogram libraries were stored. Per setup, a combined chromatogram library was created consisting of the three replicates. This performs a Percolator re-running of the combined results and provides peptide and protein quantifications at a 1% peptide and protein Q-value, respectively. For quantification, the number of minimum required and quantifiable ions were set at 5 with aligning between samples enabled.

MS2PIP, Elude and EncyclopeDIA are open source, licensed under the Apache-2.0 License, and are hosted on https://github.com/compomics/ms2pip_c, https://github.com/percolator/percolator and https://bitbucket.org/searleb/encyclopedia/wiki/Home.

### Software packages for statistics and data visualization

For basic data handling, normalization, statistics (if not stated otherwise) and annotation enrichment analysis, we used the freely available open-source bioinformatics platform Perseus (http://141.61.102.17/perseus_doku/doku.php?id=start) (v1.6.5.0). Perseus was used to visualize data from principal component analysis, non-supervised hierarchical clustering, and scatter plots. Furthermore, the 1D and 2D annotation algorithms and Fisher’s exact tests implemented in Perseus (Cox & Mann, 2012) were used for annotation enrichment analysis. Heat maps, PCA plots and bar charts were generated using GraphPad Prism software (www.graphpad.com).

### IPA Pathway analysis

For the different time points post-infection, significantly regulated human proteins (t-test, p-value ≤ 0.01; described above) were characterized by core analysis in IPA^®^ software (version 68752261, QIAGEN Inc., https://www.qiagenbioinformatics.com/products/ingenuity-pathway-analysis) using the t-test differences as directional expression values. Upstream regulators were determined using a p-value of overlap (cutoff set at 0.01) and z-scores calculated using the described algorithm implemented in IPA (69).

## Supporting information

**Figure S1.**
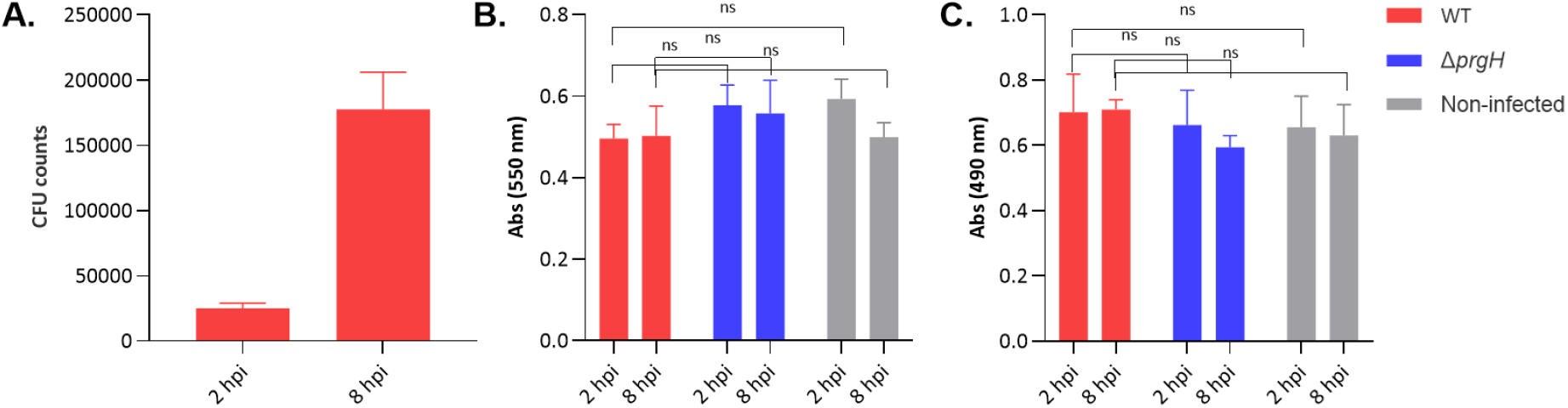
Characterization of epithelial *Salmonella* infection model. HeLa cells (1,875 x 10^4^ HeLa cells) were infected with *S*. Typhimurium at an MOI 100 and enumeration of intracellular bacteria was determined at 2 and 8 hpi by CFU counting **(A)**. Viability of *Salmonella*-infected HeLa cells was assessed in an MTT (n=4) **(B)** and an LDH assay (n=8) **(C)**. MTT reduction to formazan was measured at 595 nm and formazan formation by LDH was measured at 490 nm. Absorbance means of HeLa cells infected with *Salmonella* wild-type (WT) strain at MOI 100 at 2 and 8 hpi are not significantly different to non-infected controls and HeLa cells infected with a *ΔprgH* non-invasive control strain. Comparisons were performed using a Kruskal-Wallis’ test. Ns, not significant.

**Figure S2.**
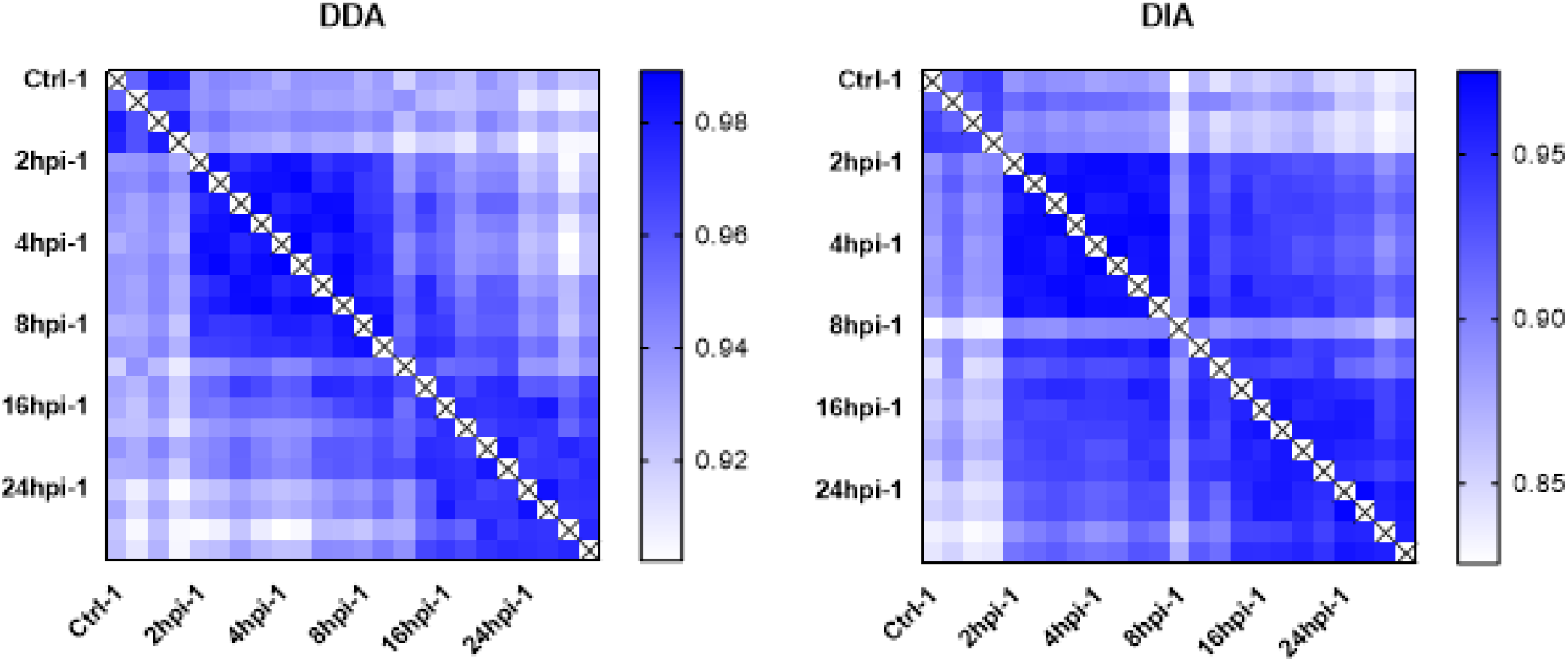
Correlations (Pearson) between all setups and replicate samples analysed. (1) Ctrl HeLa, (2) 2 hpi, (3) 4 hpi, (4) 8 hpi, (5) 16 hpi, and (6) 24 hpi.

**Figure S3.**
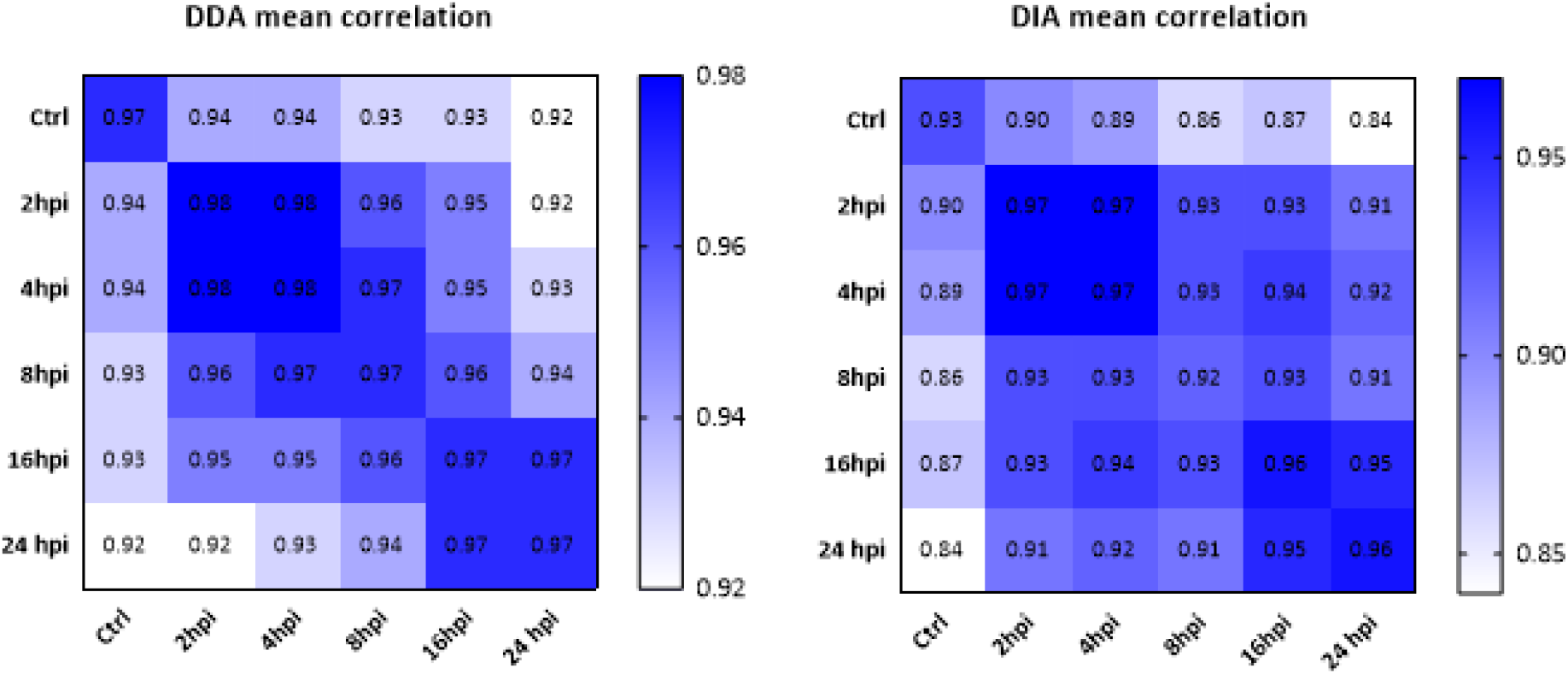
Average mean correlations (Pearson) between setups analysed. (1) Ctrl HeLa, (2) 2 hpi, (3) 4 hpi, (4) 8 hpi, (5) 16 hpi, and (6) 24 hpi.

**Figure S4.**
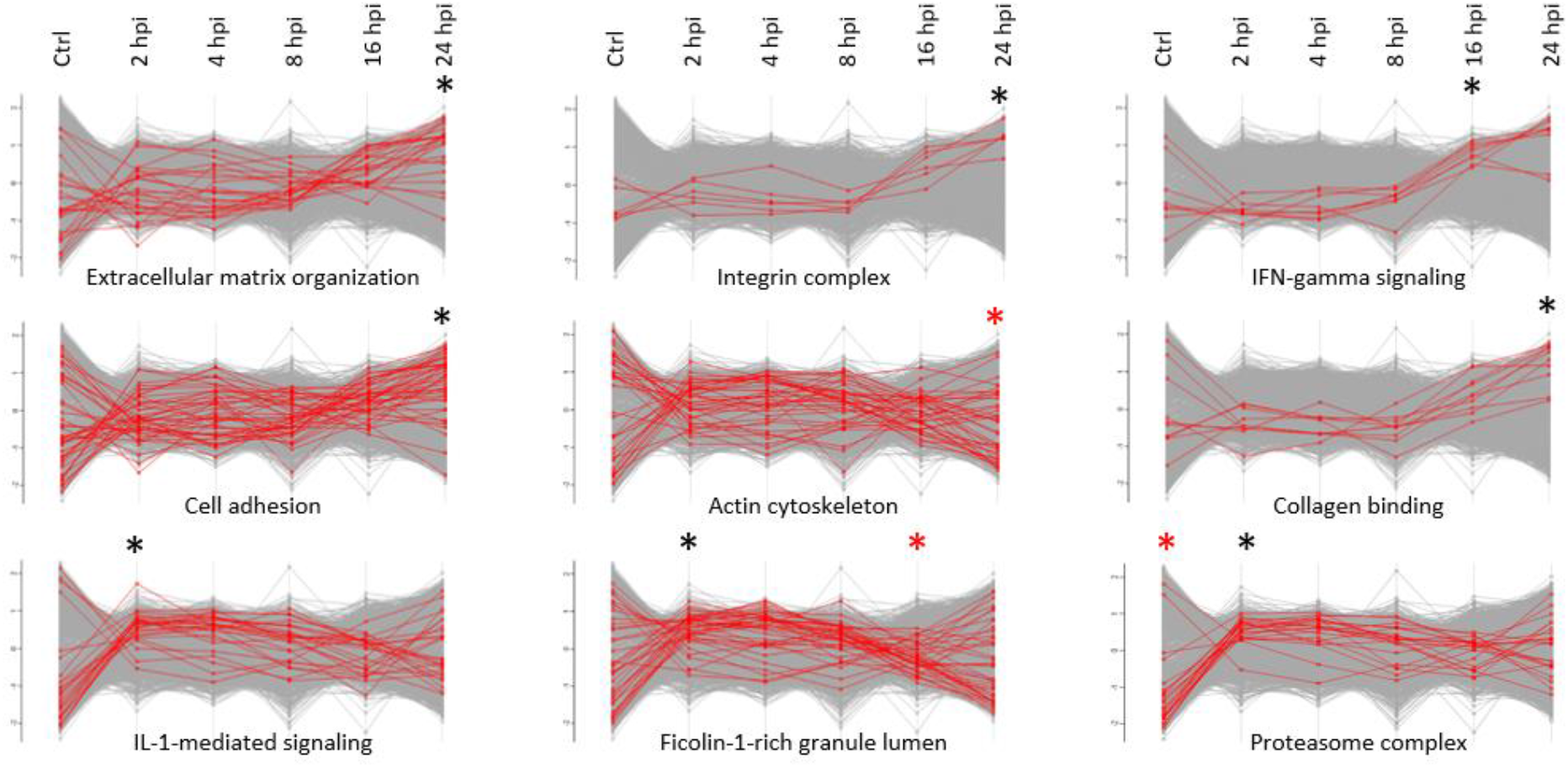
Protein profile plots of significantly regulated (FDR ≤ 0.02) normalized human annotation enrichment scores across all 6 conditions (averaged expression values) assayed by data-independent acquisition (DIA). Term enrichment was determined using the 1D annotation enrichment algorithm embedded in the Perseus software suite and p-values were corrected for multiple hypotheses testing using the Benjamini and Hochberg false discovery rate. Only corrected p-values ≤ 0.02 were considered. Selected GO terms and keyword annotations for with significantly altered protein abundancies were observed (i.e., extracellular matrix organization (GOBP, #26); upregulation at 24 hpi, cell adhesion (GOBP, #45); upregulation at 24 hpi, IL1-mediated signalling (GOBP, #27); upregulation at 2 hpi, integrin complex (keyword, #6); upregulation 24 hpi, actin cytoskeleton (GOCC, #26); downregulation 24 hpi, Ficolin-1-rich granule lumen (GOCC, #40); upregulation 2 hpi and downregulation 16 hpi, IFN-gamma signalling (GO, #8); upregulation 16 hpi, collagen binding (GO, #10); upregulation 24 hpi, proteasome complex (keyword, #22); down in control and upregulation 2 hpi) are shown for all setups analysed. Red and black asterisk above the profile plots means significant downregulation and upregulation of the corresponding proteins in the annotation group, respectively.

**Figure S5.**
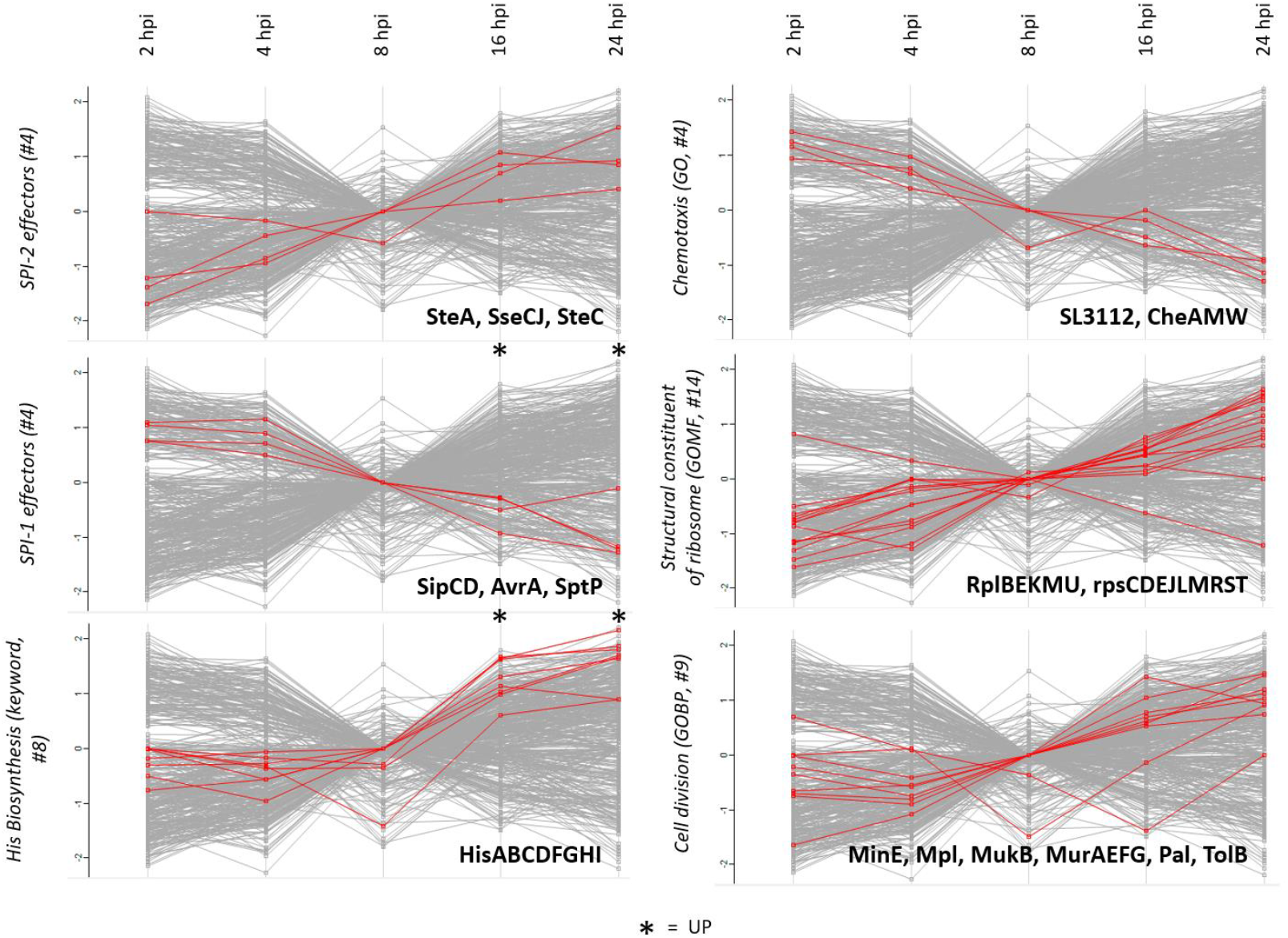
Protein profile plots of significantly regulated (FDR ≤ 0.05) normalized *S*. Typhimurium annotation enrichment scores across all 5 infection conditions (averaged expression values) assayed by data-independent acquisition (DIA). Term enrichment was determined using the 1D annotation enrichment algorithm embedded in the Perseus software suite and p-values were corrected for multiple hypotheses testing using the Benjamini and Hochberg false discovery rate. Only corrected p-values ≤ 0.05 were considered. Selected GO terms and keyword annotations for with significantly altered protein abundancies were observed (i.e., SPI-1 (#4); downregulation at 16 and 24 hpi, histidine biosynthesis (keyword, #8); upregulation at 16 and 24 hpi) are shown for the 5 infection setups analysed. A black asterisk above the profile plots means significant upregulation (by 1D annotation enrichment of the corresponding proteins in the annotation group. Besides some other categories (SPI-2 (#4), chemotaxis (GO, #4), structural constituent of the ribosome (GOMF, #14) and cell division (GOBP, #9) of which the corresponding protein members were all found to be significantly regulated based on t-testing are shown as profile plots.

## Acknowledgments

We would like to acknowledge An Staes for her help performing LC-MS sample analyses.

## Author Contributions

P.V.D. and K.G. conceived and designed the research. U.F. and P.V.D. performed the experiments. P.W. and P.V.D. analysed the data. M.D.M. performed IPA pathway analysis. M.D.M. and P.V.D. wrote the manuscript and all authors contributed to finalizing the manuscript text and gave approval to the final version of the manuscript.

## Data Availability Statement

### MS data availability

The mass spectrometry proteomics data have been deposited to the ProteomeXchange Consortium via the PRIDE (70) partner repository with the dataset identifier PXD018610 for DDA data (username, password: DwksuEy2) and with the dataset identifier PXD019863 for DIA data (username, password: CRmowDjV).

The generated spectral libraries, FASTA databases, and search results are organized in the Open Science Framework (OSF) project page (71) and are available at https://osf.io/u96bk/.

## Funding

P.V.D. acknowledges funding from the European Research Council (ERC) under the European Union’s Horizon 2020 research and innovation program (PROPHECY grant agreement No 803972) and support from the Research Foundation – Flanders (FWO-Vlaanderen), project number G051120N. K.G. acknowledges support from a Ghent University Concerted Research Actions (grant BOF14/GOA/013).

## Competing interests

The authors have declared that no competing interests exist.

## Supporting information Supplementary Table captions

**Table S1A | List of 6,696 proteins identified in the dual proteome shotgun samples of *S*. Typhimurium infected human HeLa cells by means of data-dependent acquisition (DDA).** *S*. Typhimurium infected human HeLa cells were prepared to monitor quantitative differences in steady-state protein expression levels at 2, 4, 8, 16 and 24 hpi in biological quadruplicates. UniProt database primary accession number, UniProt entry (name), gene name, corresponding protein description and sequence, molecular protein weight, number of (razor/unique) peptides identified, (razor/unique) sequence coverage, score, log2-transformed LFQ protein intensities, GO annotation and keywords, MS/MS counts and sequence length(s) are given. Ranking is done alphabetically according to protein names.

**Table S1B | List of 6,734 proteins identified in the dual proteome shotgun samples of *S*. Typhimurium infected human HeLa cells by means of data-independent acquisition (DIA).** *S*. Typhimurium infected human HeLa cells were prepared to monitor quantitative differences in steady-state protein expression levels at 2, 4, 8, 16 and 24 hpi in biological quadruplicates. UniProt database primary accession number, UniProt entry (name), gene name, corresponding protein description and sequence, sequence length, number (sequence) of peptides identified, species, log2-transformed protein intensities, GO annotation and keywords are given. Ranking is done alphabetically according to protein names.

**Table S2A| List of 4,666 human proteins quantified with a minimum of three valid values in at least one setup of the dual proteome shotgun samples of *S*. Typhimurium infected human or control HeLa cells by means of data-dependent acquisition (DDA).** Comparisons were analysed by *t* testing and significant hits determined using as cut-off values a permutation based false discovery rate (FDR) of 0.01 and 0.001 (100 permutations) and a background variance parameter S0 of 0.1. Table headers are as in Supplementary Table S1A.

**Table S2B| List of 479 *S*. Typhimurium proteins quantified with a minimum of three valid values in at least one setup of the dual proteome shotgun samples of *S*. Typhimurium infected human HeLa cells by means of data-dependent acquisition (DDA).** Comparisons were analysed by *t* testing and significant hits determined using as cut-off values a permutation based false discovery rate (FDR) of 0.05 and 0.01 (100 permutations) and a background variance parameter S0 of 1. Table headers are as in Supplementary Table S1A and cluster indication is as in Figure 2 (DDA data).

**Table S2C| List of 5,818 human proteins quantified with a minimum of three valid values in at least one setup of the dual proteome shotgun samples of *S*. Typhimurium infected human or control HeLa cells by means of data-independent acquisition (DIA).** Comparisons were analysed by *t* testing and significant hits determined using as cut-off values a permutation based false discovery rate (FDR) of 0.01 and 0.001 (100 permutations) and a background variance parameter S0 of 0.1. Table headers are as in Supplementary Table S1B and cluster indication is as in Figure 3.

**Table S2D | List of 844 *S*. Typhimurium proteins quantified with a minimum of three valid values in at least one setup of the dual proteome shotgun samples of *S*. Typhimurium infected human HeLa cells by means of data-independent acquisition (DIA).** Comparisons were analysed by *t* testing and significant hits determined using as cut-off values a permutation based false discovery rate (FDR) of 0.05 and 0.01 (100 permutations) and a background variance parameter S0 of 1. Table headers are as in Supplementary Table S1B with regulon and genome locus and strand info added, and cluster indication is as in Figure 2 (DIA data).

**Table S3| Significantly regulated (FDR ≤ 0.02) normalized human annotation enrichment scores across all 6 conditions (averaged expression values) assayed by data-independent acquisition (DIA)**. Term enrichment was determined using the 1D annotation enrichment algorithm embedded in the Perseus software suite and p-values were corrected for multiple hypotheses testing using the Benjamini and Hochberg false discovery rate.

